# Maternal glucocorticoids promote offspring growth without inducing oxidative stress or shortening telomeres in wild red squirrels

**DOI:** 10.1101/680421

**Authors:** Ben Dantzer, Freya van Kesteren, Sarah E. Westrick, Stan Boutin, Andrew G. McAdam, Jeffrey E. Lane, Robert Gillespie, Ariana Majer, Mark Haussmann, Pat Monaghan

## Abstract

Elevations in glucocorticoid levels (GCs) in breeding females may induce adaptive shifts in offspring life histories. Offspring produced by mothers with elevated GCs may be better prepared to face harsh environments where a faster pace of life is beneficial. We examined how experimentally elevated GCs in pregnant or lactating North American red squirrels (*Tamiasciurus hudsonicus*) affected offspring postnatal growth, structural size, oxidative stress levels (two antioxidants and oxidative protein damage) in three different tissues (blood, heart, liver), and liver telomere lengths. We predicted that offspring from mothers treated with GCs would grow faster but would also have higher levels of oxidative stress and shorter telomeres, which may predict reduced longevity. Offspring from mothers treated with GCs during pregnancy were 8.3% lighter around birth but grew (in body mass) 17.0% faster than those from controls, whereas offspring from mothers treated with GCs during lactation grew 34.8% slower than those from controls and did not differ in body mass around birth. Treating mothers with GCs during pregnancy or lactation did not alter the oxidative stress levels or telomere lengths of their offspring. Fast-growing offspring from any of the treatment groups did not have higher oxidative stress levels or shorter telomere lengths, indicating that offspring that grew faster early in life did not exhibit oxidative costs after this period of growth. Our results indicate that elevations in maternal GCs may induce plasticity in offspring growth without long-term oxidative costs to the offspring that might result in a shortened lifespan.

**Summary Statement:** We show that experimental increases in glucocorticoids in breeding female North American red squirrels affects offspring postnatal growth but not levels of oxidative damage and antioxidants or telomere lengths.

## Introduction

Parents can have long-lasting impacts on their offspring across a diversity of taxa. These parental effects have drawn substantial interest because they suggest that parental characteristics or the parental environment itself could induce adaptive shifts in offspring traits that prepare them for specific environments (i.e., adaptive transgenerational phenotypic plasticity: Mousseau and Fox, 1998; Agrawal et al., 1999; Galloway and Etterson, 2007; Dantzer et al., 2013). Furthermore, changes in maternal hormone levels, especially glucocorticoids (GCs), are widely suspected to act as a mediator of transgenerational phenotypic plasticity in vertebrates (Love and Williams, 2008; Sheriff et al., 2017).

GCs are metabolic hormones released by the hypothalamic-pituitary-adrenal (HPA) axis (Sapolsky et al., 2000), in response to a variety of ecologically salient cues. In mammals, studies in laboratory animals (Barbazanges et al., 1996; Harris and Seckl, 2011), 10) and in humans (Weinstock, 2005; Lupien et al., 2009) show that elevated maternal GCs can generate stable individual differences in offspring physiology and behaviour through the transfer of maternally-derived GCs to offspring across the placenta (Barbazanges et al., 1996; Lupien et al., 2009), in milk (Zarrow et al., 1970; Casolini et al., 1997), or through changes in maternal behaviour (Herrenkohl and Whitney, 1976; Brummelte and Galea, 2010; Nephew and Bridges, 2011). Maternal GCs could also act as an internal cue for offspring to modify their own development (Nettle et al., 2013). Regardless of the pathway, there is much evidence that changes in maternal GCs mediate parental effects whose influence may even persist across generations via epigenetic mechanisms (Weaver et al., 2004).

Some of these changes in offspring characteristics that are caused by elevations in maternal GCs are suspected to reflect adaptive plasticity in offspring life history traits, such as modifying the trade-off between early life growth and lifespan (Monaghan and Haussmann, 2015; Berghänel et al., 2017). Elevated maternal GCs may induce a “faster” life history strategy whereby offspring produced by mothers with elevated GCs grow or develop faster (Dantzer et al., 2013; Berghänel et al., 2016). Such adjustments in the “pace of life” may be adaptive, as fast postnatal growth or a quicker developmental time may be beneficial when the risk of extrinsic mortality is heightened (Reznick et al., 1990; Stearns et al., 2000), which may coincide with elevated maternal GCs. However, this increased investment in postnatal growth is expected to carry costs for offspring longevity whereby fast-growing individuals exhibit a shortened lifespan (Metcalfe and Monaghan, 2001; Lee et al., 2013; Lemaître et al., 2015; Monaghan and Ozanne, 2018).

Identifying if maternal stress induces a trade-off between growth and lifespan in wild animals is needed, as any costs of increased exposure to maternal GCs may be masked by benign environmental conditions in the laboratory. Accurately documenting lifespan in wild animals remains challenging in many species, and the stochastic nature of mortality in wild animals may obscure any mechanistic costs of increased exposure to maternal GCs for offspring. Consequently, one way to examine if elevated maternal GCs induce a trade-off between growth and longevity in offspring is to examine how it affects the possible underlying mechanisms of reduced longevity or physiological correlates that may predict a shortened lifespan. The free-radical theory of aging (Beckman and Ames, 1998) provides one framework to examine the mechanisms by which maternal GCs may induce this trade-off between offspring growth and longevity. Reactive oxygen species (ROS) produced during aerobic respiration can have damaging effects on cells (Beckman and Ames, 1998; Halliwell and Gutteridge, 2015). ROS production may be elevated by increased aerobic respiration due to enhanced investment in growth or reproduction (Fletcher et al., 2012; Blount et al., 2016; Smith et al., 2016; Monaghan and Ozanne, 2018) or by increased GCs (Kotrschal et al., 2007; You et al., 2009; Costantini et al., 2011; Herborn et al., 2014). Antioxidants produced by individuals (enzymatic antioxidants such as superoxide dismutase) as well as those antioxidants from the external environment (non-enzymatic antioxidants in the diet) can lessen the impact of ROS production (Selman et al., 2012). An important type of oxidative damage occurs to the protective ends of chromosomes, called telomeres. Telomeres are the repetitive DNA sequences that occur at the ends of eukaryote chromosomes whose length is shortened during each cell division (Harley et al., 1990; Aubert and Lansdorp, 2008), and may also be reduced by the increased production of ROS (von Zglinicki, 2002; Houben et al., 2008; Reichert and Stier, 2017; Monaghan and Ozanne, 2018). Cells with a reduced telomere length become senescent, stop dividing, and undergo apoptosis unless the enzyme telomerase, or another elongation process, is produced to elongate the telomeres (Bodnar et al., 1998; Rudolph et al., 1999). Telomere length or rate of loss has been found to be predictive of the mortality risk of individuals (Cawthon et al., 2003; Haussmann et al., 2005; Bize et al., 2009; Salomons et al., 2009; Boonekamp et al., 2014; Munoz-Lorente et al., 2019), though the strength of this relationship may vary among taxa and in relation to other life history traits. Furthermore, avian or mammalian species with longer lifespans have been shown to exhibit slower age-specific rates of telomere loss (Haussmann et al., 2003; Dantzer and Fletcher, 2015; Sudyka et al., 2016; Tricola et al., 2018).

Elevated exposure to maternal GCs may, therefore, induce a life history trade-off between offspring growth and longevity, because offspring may experience elevated oxidative damage, decreased antioxidant levels, and/or shortened telomeres, either due to oxidative stress or increased cell division associated with elevated growth (Shalev, 2012; Haussmann and Heidinger, 2015; Monaghan and Haussmann, 2015; Shalev and Belsky, 2016; Monaghan and Haussmann, 2018). Previous studies across taxa show that elevated exposure to maternal GCs can shorten telomere lengths in offspring (Entringer et al., 2011; Haussmann et al., 2012; Herborn et al., 2014; Marchetto et al., 2016) or increase their rate of attrition as they age (Haussmann and Heidinger, 2015), which could cause or be associated with a shortened lifespan. For example, experimental studies in captive and wild birds show that offspring that had exogenous GCs added to their eggs or were administered GCs during chick growth had a heightened physiological stress response, higher levels of oxidative stress, and shorter telomeres early in life (Haussmann et al., 2012; Herborn et al., 2014). Despite much interest in this topic, few studies in wild animals have examined if experimental elevations in the GCs of breeding females impact the oxidative state of offspring or explicitly tested the prediction that elevations in the GCs of breeding females increase early life growth. Additionally, few studies have tested whether elevated GCs in breeding females or fast early life growth comes at some cost by promoting oxidative stress and shortening telomeres in offspring.

We tested the hypothesis that elevations in maternal GCs would promote a faster life history strategy in offspring of wild North American red squirrels (*Tamiasciurus hudsonicus*). We treated females with GCs using a protocol that allowed us to increase circulating GCs within a physiologically-relevant range (van Kesteren et al., 2019). We treated females with GCs either during pregnancy or lactation to assess if the timing of exposure to maternal GCs influenced their effects on offspring. Other than for offspring growth in body mass, we did not have strong *a priori* expectations of how the timing of elevated maternal GCs would differentially impact offspring because elevated maternal GCs during pregnancy or lactation can impact offspring through the same pathways: direct transfer of maternal GCs to offspring across the placenta or through milk (“programming”), altering maternal behaviour, or affecting offspring behaviour (see references above). However, based upon our previous study (Dantzer et al., 2013), we predicted that offspring produced by mothers treated with GCs during pregnancy would grow faster in body mass. We did not have an *a priori* expectation for how treating mothers with GCs during lactation would impact offspring growth in body mass, though results from a previous study suggested that it should reduce growth (Nephew and Bridges, 2011).

We measured offspring postnatal growth in body mass prior to weaning (∼1 to 25 d of age) and subsequently obtained measures of oxidative stress when pups were weaned (∼70 d of age). In three tissues (liver, heart muscle, and blood) collected from weaned offspring, we measured one enzymatic antioxidant (superoxide dismutase), one type of non-enzymatic antioxidant (total antioxidant capacity), and one type of oxidative damage (protein damage measured via protein carbonyls). We used multiple tissues because other studies have highlighted how experimental manipulations can have tissue-specific effects (Garratt et al., 2012). To assess the cumulative impact of elevated maternal GCs on the oxidative state of offspring and how offspring growth impacted telomere lengths, we also measured telomere lengths in DNA from the liver. We focused on liver telomere lengths because the liver is a mitotically active tissue, produces growth hormones, and previous studies have documented a reduction in telomere length with faster growth (Monaghan and Ozanne, 2018), suggesting that the liver would be a good tissue to investigate any oxidative costs of growth. Although we only measured telomere lengths in one tissue, previous studies indicate that telomere lengths measured in one somatic tissue are strongly correlated with those in others (Friedrich et al., 2000). Because our estimate of offspring growth from ∼1 to 25 d of age was temporally separated from when we obtained our measures of oxidative stress and telomere lengths (when offspring were weaned [∼70 d]), we were able to assess whether there were persistent or cumulative oxidative costs to fast growth.

We predicted that offspring from mothers treated with GCs during pregnancy would grow quicker in body mass after birth, but would experience more oxidative stress (manifested as a reduction in antioxidants and an increase in oxidative damage) and decreased telomere length, which would be a result of increased oxidative stress or increased cell division associated with faster growth. Because we have previously found that female red squirrels can ameliorate the trade-off between offspring number and growth (Dantzer et al., 2013; Westrick et al., 2019 preprint), we examined if elevated maternal GCs altered the trade-off between litter size and offspring growth or structural (skeletal) size. Previous studies highlight that elevated maternal GCs can impact offspring birth weight (Berghänel et al., 2017) so we also examined the treatment effects on the first measure of body mass. Because early life exposure to GCs may modify the direction and strength of the association between two variables (Careau et al., 2014; Merill and Grindstaff, 2018), we also examined if increases in maternal GCs affected the expected negative relationship between offspring growth and oxidative stress state (Smith et al., 2016) by assessing the statistical interaction between offspring growth and maternal treatment.

## Materials and Methods

### Study area & measuring offspring growth

We conducted this study as a part of a long-term study of red squirrels in the Yukon, Canada that takes place on the traditional territory of the Champagne and Aishihik First Nations. Squirrels in our study population were all marked individually with unique ear tags and combinations of coloured wire threaded through the ear tags (McAdam et al., 2007). Females in our study population usually produce one litter in the spring, and rarely produce more than one litter of offspring to weaning per year (Boutin et al., 2006). Adult females were captured and handled every ∼3 to 21 d to assess reproductive status through abdominal palpation and nipple condition. Pups were retrieved from the nest two times. The first nest entry occurred immediately after parturition and the second nest entry occurred when pups were approximately 25 d of age. At both nest entries, pups were briefly removed from their nest, sexed, and weighed to the nearest 0.01 g using a portable balance. At the first nest entry, we marked them uniquely by obtaining a small ear biopsy (for later paternity analyses) and then we permanently marked pups at the second nest entry with unique metal ear tags. Because offspring growth in body mass during this period of time is approximately linear (McAdam et al., 2002), we estimated offspring growth as the change in body mass from the first to second nest entry divided by the total number of days elapsed between the two measures of body mass. Red squirrels usually first emerge from their nest around 30-35 d old and are usually weaned around 70 d old, such that this period of growth from ∼1-25 d of age represents a period in which offspring are only consuming milk from their mother. At the second nest entry, we measured zygomatic arch width and right hind foot length to the nearest 1 mm using digital callipers or a ruler, respectively. Because we only obtained these morphological measures at the second nest entry (when pups were ∼25 d old), we were not able to measure the change in offspring structural size as we did for growth. All of our procedures were approved by the Animal Care and Use Committee at the University of Michigan (PRO00007805).

### Maternal treatments

We used four separate treatment groups to assess the effects of elevated maternal GCs on offspring over four different years (2012, 2015-2017), although in 2012 we only collected growth in body mass data. Individual adult females were treated with GCs either during pregnancy or lactation (“Pregnancy GCs” and “Lactation GCs”), whereas other females were treated as controls during pregnancy or lactation (“Pregnancy Controls” or “Lactation Controls”). We increased maternal GCs either during pregnancy or lactation using an established experimental protocol (Dantzer et al., 2013; van Kesteren et al., 2019). Briefly, we treated females in the Pregnancy GCs (n = 44 total litters from 43 unique females) and Lactation GCs (n = 18 litters from 17 unique females) treatment groups with exogenous cortisol (hydrocortisone, Sigma H4001) dissolved in peanut butter and wheat germ mixture (8 g of peanut butter, 2 g of wheat germ). Females in the Pregnancy Control (n = 31 total litters from 32 unique females) or Lactation Control (n = 17 litters from 16 unique females) treatment groups were fed the same amount of peanut butter and wheat germ mixture but lacking the hydrocortisone. GC treatments were prepared by dissolving hydrocortisone in 1 mL of 100% ethanol and then 5 mL of 100% peanut oil before allowing the emulsion to sit overnight so that the ethanol could evaporate. The following morning, the hydrocortisone emulsion was thoroughly mixed with the appropriate amount of peanut butter and wheat germ, weighed out into individual dosages (∼10 g each), placed into an individual container, and then frozen at −20 °C until provisioning to the treated squirrels.

Each day during the treatment period, we placed individual dosages into a bucket that was hung ∼7-10 m off the ground on the centre of the squirrel’s territory. Squirrels defend these buckets from all other conspecifics and heterospecifics (van Kesteren et al., 2019), so we can, therefore, be confident that the squirrels that were given these treatments were consuming them. The dosage of hydrocortisone varied among some of the treatment groups but we have previously shown that either 3, 6, 8, or 12 mg of hydrocortisone per day significantly elevates baseline plasma cortisol and faecal glucocorticoid metabolite levels but within a physiologically-relevant range (Dantzer et al., 2013; van Kesteren et al., 2019). Females in the Pregnancy GCs treatment group were provisioned either with 3 mg (n = 4 litters), 6 mg (n = 6 litters), 8 mg (n = 26 litters), or 12 mg (9 litters) of hydrocortisone whereas females in the Lactation GCs treatment groups were provisioned either with 8 mg per day (n = 10 litters) or 12 mg per day (n = 8 litters). Although the dosage administered to females treated during pregnancy or lactation varied, we grouped those administered GCs in the same treatment group regardless of dosage (i.e., Pregnancy GCs contained females administered 3-12 mg of hydrocortisone per day, Lactation GCs contained females administered 8 or 12 mg of hydrocortisone per day). We did this for three reasons. First, we have previously shown that females provisioned with 6, 8, or 12 mg of hydrocortisone per day did not differ in their faecal glucocorticoid metabolite levels (FGM), although there were non-significant trends where squirrels fed higher dosages of hydrocortisone had higher FGM (van Kesteren et al., 2019). Second, in preliminary analyses of the data presented here and in a previous study (Dantzer et al., 2013), we found that the effects of different GC dosages (3, 6, 8, and 12 mg per day) that were provided to pregnant females on offspring growth were in the same direction (increased postnatal growth: Table S1). For females treated with Lactation GCs (8 or 12 mg per day), we also found that the effects on offspring growth were in the same direction (decreased postnatal growth: Table S1). Finally, our sample sizes in some of the treatment groups (3 and 6 mg per day in the Pregnancy GCs treatment group) were too small to assess whether there were statistical differences among the different dosage groups.

We aimed to treat females in the pregnancy treatments from approximately 20 d after conception until 5 d after birth (or 20-40 d post-conception as red squirrels on average have a ∼35 d gestation period), whereas we actually treated females in the pregnancy GCs treatment group from 24.2 ± 0.9 d (mean ± SE) to 39.0 ± 0.6 d post-conception (mean ± SE treatment duration: 14.9 d ± 0.8 d) and females in the pregnancy control treatment group from 20.4 ± 0.8 d to 38.6 ± 0.5 d (treatment duration: 18.2 ± 0.8 d). We aimed to treat females in the lactation treatments from approximately 5 d after parturition until 15 d post-parturition, whereas we actually treated females in the Lactation GCs treatment group from 5.4 ± 0.5 d (mean ± SE) to 14.5 ± 0.6 d post-conception (mean ± SE treatment duration: 9.1 d ± 0.1 d) and females in the Lactation Control treatment group from 4.9 ± 0.4 d to 14.1 ± 0.5 d (treatment duration: 9.2 ± 0.5 d). Given a ∼35 d gestation period and a ∼70 d lactation period in this population, our pregnancy treatments corresponded to treating females during the last trimester of gestation and into the first few days of lactation, whereas our lactation treatments corresponded to early lactation before offspring begin to feed independently (they typically leave the nest on their own for the first time at ∼35 d). Note that this means that the lactation treatments occurred during a time when the offspring would not be able to consume any of the treatments themselves so any effects on offspring were likely due to the maternal phenotype.

Some females in the Pregnancy GCs (n = 3 litters) or Pregnancy Control (n = 2 litters) treatment groups aborted their litters prior to the first nest entry (no females in the lactation treatments aborted their litters prior to the first nest entry). Some females lost their litters (likely due to nest predation: Studd et al., 2015) after the first nest entry but before the second nest entry when we could obtain the second measure of pup body mass to estimate their growth and the only measures of offspring morphology, thereby reducing our sample sizes to estimate the treatment effects on the first measure of offspring body mass, postnatal growth and body size (sample sizes shown in Tables 2-4, S1-S5). Below we show that there was no evidence that the treatments had differential effects on litter survival from the first to second nest entry (see Results).

### Tissue sample collection

Pups are weaned when they are approximately ∼70 d of age and generally stay on their natal territory until dispersal soon after (Larsen and Boutin, 1994). When juvenile squirrels were ∼70 d of age, they were euthanized and tissues (liver and cardiac muscle) were immediately removed, rinsed with PBS buffer, snap frozen on dry ice, and then stored in liquid nitrogen or in a −80 °C freezer until analysis. Trunk blood was collected through decapitation and then centrifuged at 10,000 *g* for 10 min at room temperature to separate plasma and red blood cells.

### Haematocrit

We measured packed red blood cell volume (haematocrit) as a measure of body condition as some (though not all) previous studies have found that higher haematocrit levels correspond to better body condition or improved reproductive performance, at least in some studies of wild birds (Breuner et al., 2013; Minias, 2015; Fronstin et al., 2016). Before pups were euthanized, we collected a blood sample from the hind nail into a heparinized capillary tube. Haematocrit was quantified using a micro-capillary reader after centrifuging blood samples at 10,000 *g* for 10 min at room temperature.

### Protein carbonyls

We measured oxidative damage to proteins (Monaghan et al., 2009) using the protein carbonyl colorimetric kit by Cayman Chemical (Ann Arbor, USA). Briefly, ∼200 mg of cardiac muscle or liver were homogenized in ∼1000 µL of 50mM MES buffer containing 1mM EDTA using a sonicator, and then centrifuged at 10,000 *g* for 15 min at 4 °C. The protein concentration of tissue homogenate supernatant and plasma samples was measured prior to the assay using a Biotek Take3 protocol (Biotek, Vermont, USA) and samples were diluted in PBS buffer to give a protein range between 1-10 mg/ml, as recommended by the manufacturer. The average intra-assay CV for samples for plasma, heart, or liver were 1.2%, 2.8%, and 1.6%, respectively. Inter-assay CVs for a red squirrel pooled sample run on repeat assays for plasma (n = 7 assays), heart (n = 6), or liver (n = 7) were 3.5%, 9.0%, and 8.3%, respectively. We also ran a positive control (oxidized bovine serum albumin) in two different assays and the inter-assay CV was 3.1%.

### Superoxide dismutase

We obtained one measure of the levels of enzymatic antioxidants (Monaghan et al., 2009) by quantifying levels of superoxide dismutase (SOD) using the SOD kit from Cayman Chemical. SOD was expressed as units/mg/ml protein (quantified using a Biotek Take3 protocol). Red blood cells were lysed as per the manufacturer’s protocol. The average intra-assay CV for samples for RBCs, heart, or liver were 2.3%, 4.7%, and 3.1%, respectively. Inter-assay CVs for a red squirrel pooled sample run on repeat assays for RBCs (n = 10 assays), heart (n = 2), or liver (n = 4) were 16.9%, 4.6%, and 6.5%, respectively.

### Total antioxidant capacity

We obtained one measure of the levels of non-enzymatic antioxidants (Monaghan et al., 2009) by quantifying total antioxidant capacity (TAC) using the TAC kit from Cayman Chemical. Plasma was diluted in assay buffer and assayed according to the manufacturer’s protocol. Liver and cardiac muscle (∼47 mg) were separately homogenized in 250 µL PBS using a sonicator and the supernatant was diluted in assay buffer and used in the assay. The average intra-assay CVs for samples for plasma, heart, or liver were 4.7%, 3.2%, and 4.7%, respectively. The inter-assay CVs for standards run on all the plates for plasma (n = 5 assays) was 15.8% whereas the inter-assay CV for a red squirrel pooled sample run on repeat for heart (n = 8) or liver (n = 2) were 15.4% and 5.4%, respectively.

### Telomeres

Liver telomere lengths were measured using the telomere restriction fragment (TRF) assay following established methods (Haussmann and Mauck, 2008). Briefly, 2 to 10 g slices of liver tissue were homogenized in cell lysis solution and proteinase K (Qiagen, Germantown, USA). DNA was extracted from the liver homogenates and resuspended in buffer. The resuspended DNA was restriction digested with 15 U of HinfI, 75U of HaeIII and 40U of RsaI (New England BioLabs, Ipswich, USA) at 37 °C. DNA was then separated using pulsed field electrophoresis at 14°C for 19 hours followed by in-gel hybridization overnight at 37°C with a radioactively labeled telomere-specific oligo (CCCTAA)_4_. Hybridized gels were placed on a phosphorscreen (Amersham Biosciences, Buckinghamshire, UK), which was scanned on a Typhoon Imager (Amersham Biosciences). Densitometry in ImageJ (v. 1.51s) was used to determine the position and the strength of the radioactive signal in each of the lanes compared with the molecular marker (Quick-Load1 kb DNA Extend DNA Ladder; New England BioLabs) to calculate telomere lengths for each sample. Inter-gel variation was accounted for by calculating the mean TRF length of standard samples run on each gel.

### Statistical analyses

We assessed the effects of maternal treatments on litter survival from the first to second nest entry (proportion of pups present at both first and second nest entries) and litter size and litter sex ratio (proportion of litter composed of males) as recorded at the first and second nest entry using generalized linear mixed-effects models (GLMMs: litter survival and litter sex ratio, using binomial errors) or a linear mixed-effect model (LMM: litter size). Each of these separate models contained maternal treatment, birth date, and year as fixed effects. Models for lactation and pregnancy treatments were run separately. There was one litter from a Pregnancy Control treatment female where her litter size at the second nest entry was greater than the at the first, likely because we missed a pup in the nest at the first nest entry, and we therefore excluded this litter from our analyses of the effects of the treatments on litter survival, litter size, and litter sex ratio. We confirmed that none of the GLMMs were overdispersed as all dispersion parameters were <1.

We assessed the effects of maternal treatments on offspring body mass at the first nest entry soon after parturition (birth mass), growth in postnatal body mass, and a single measure of size using separate LMMs for pregnancy and lactation treatments. Each of the six LMMs (one model for birth mass, one model for growth, and one model for size for pregnancy treatments; one model for birth mass, one model for growth, and one model for size for lactation treatments) included a fixed effect for maternal treatment and covariates (sex, year [categorical variable], birth date, litter size) that could impact offspring birth mass, growth, or size. We included a two-way interaction term between treatment and litter size to identify if elevations in maternal GCs altered the trade-off between litter size and offspring birth mass, growth, or size, as shown previously for offspring growth (Dantzer et al., 2013). We included a two-way interaction between treatment and sex to assess if the treatments had sex-specific effects on birth mass, growth, and size, as documented in other species (Dantzer et al., 2019). In our model for the effects of the treatments on offspring birth mass, we included a fixed effect for age of the pups to control for the impact of age on body mass. We used a principal component analysis (PCA) using a covariance matrix in the R package ade4 (version 1.7-13, Dray and Dufour, 2007) to generate a composite score of offspring size. The first principal component axis (PC1, hereafter “size”) explained 69.9% of the variation in offspring size as measured by zygomatic arch width and hind foot length. Both zygomatic arch width (0.71) and hind foot length (0.71) loaded positively on PC1, indicating that larger PC1 scores corresponded to offspring with longer hind feet and wider crania.

Oxidative stress reflects an imbalance between antioxidants and the production of ROS that can damage proteins, lipids, or DNA (Monaghan et al., 2009). Consequently, the effects of our treatments on measures of antioxidants should not be viewed in absence of their effects on our measures of oxidative damage (Costantini and Verhulst, 2009). We used a PCA to create a composite variable that reflected the oxidative state of an offspring. The PCA was composed of the two antioxidants (SOD, TAC) and one measure of oxidative damage (PCC). We conducted a separate PCA (using a correlation matrix) for each tissue type using the package ade4. For some individuals, we were missing measures of TAC (heart: n = 3; plasma: n = 2) or PCC (heart: n = 2; plasma: n = 4) so we substituted average values for the PCA.

Low scores for Blood PC2, Heart PC2, or Liver PC1 corresponded to squirrels that were exhibiting oxidative stress as they represented lower levels of the two antioxidants (SOD, TAC) for blood and liver tissue or just one antioxidant (SOD) for heart tissue and, for heart tissue, higher levels of protein damage (PCC, Table 1). We used these composite variables describing oxidative state of each tissue, haematocrit, or telomere length as the response variables in separate LMMs. Each of these LMMs contained a fixed effect for maternal treatment and offspring sex, year [categorical variable], birth date, and litter size. Because offspring growth may impact oxidative stress levels or telomere lengths (Monaghan and Ozanne, 2018), we included a two-way interaction term between treatment and offspring growth to examine if mothers with elevated maternal GCs exhibited an altered relationship between growth and the response variable (Careau et al., 2014; Merill and Grindstaff, 2018). Due to smaller sample sizes for these variables, we did not include an interaction between sex and treatment. In the model to assess treatment effects on telomere lengths, we also included a fixed effect for the oxidative stress levels in the liver (Liver PC1).

**Table 1.**
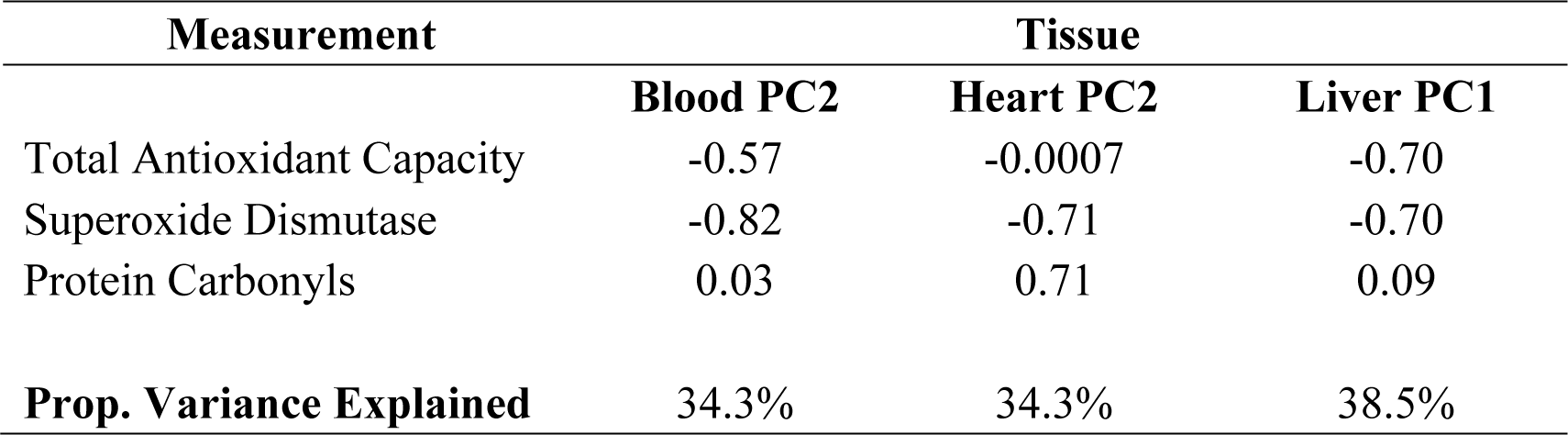
Results from principal component analyses to derive axes of variation of oxidative stress state in weaned pups. For each principal component shown, high values correspond to lower oxidative stress levels as they reflect samples with low levels of two (Blood PC2, Liver PC1) or one (Heart PC2) antioxidants (total antioxidant capacity, superoxide dismutase) and for heart tissue, higher levels of oxidative protein damage (protein carbonyls).

| Measurement | Tissue |  |  |
| --- | --- | --- | --- |
|  | Blood PC2 | Heart PC2 | Liver PC1 |
| Total Antioxidant Capacity | -0.57 | -0.0007 | -0.70 |
| Superoxide Dismutase | -0.82 | -0.71 | -0.70 |
| Protein Carbonyls | 0.03 | 0.71 | 0.09 |
| **Prop. Variance Explained** | 34.3% | 34.3% | 38.5% |

All analyses were conducted in R (version 3.5.2, R Development Core Team, 2013) using lme4 (version 1.1-18-1, Bates et al., 2015) and *P*-values for fixed effects were estimated using lmerTest (version 3.0-1, Kuznetsova et al., 2017). Continuous predictor variables were standardized (mean of 0, SD of 1) with birth date, litter size, and growth being standardized within each grid-year-treatment combination. Assumptions of homoscedasticity, normality of residuals for our LMMs, and a lack of high leverage observations were confirmed using diagnostic plots (Zuur et al., 2010). As we only had one estimate of birth mass, growth, size, oxidative stress, or telomere lengths per individual pup, we did not include random intercept terms for individual identity in any of the models described above. Whenever we had repeated observations of the same litter (e.g., multiple estimates of offspring growth from different pups within the same litter), we included a random intercept term for litter identity. However, we did not include a random intercept in some models (shown in electronic supplementary materials) due to model convergence issues that were likely the result of having only a few observations of multiple pups being measured within one litter or for having a relatively small number of litters. As we had few observations of the same females across years for most of our response variables (sample sizes shown above), we did not include a random intercept for maternal identity as this was usually redundant with litter identity. Random effects were not included in our models of the effects of the treatments on litter characteristics (survival, size, sex ratio) as we only had a few females observed across multiple years. We estimated variance inflation factors (VIFs) from our models to assess multicollinearity among the predictor variables (Zuur et al., 2010) and VIFs indicated that multicollinearity was not an issue in these models (all VIF < 3 except if included in an interaction or a multi-level categorical variable).

## Results

### Effects of treatments on litter survival, litter size, & litter sex ratio

There was no evidence that treating mothers with GCs during pregnancy or lactation caused litter failure or altered litter size or litter sex ratio compared to the controls. For those mothers producing offspring until at least the first nest entry (occurring soon after pups were born), the proportion of the litter that survived from the first to the second nest entry did not differ between mothers treated with GCs during pregnancy (n = 42 litters, 53.8 ± 7% of total pups survived from the first to second nest entry) and the pregnancy (time-matched) controls (n = 29 litters, 63 ± 8%, *z* = −0.66, *P* = 0.51), nor between mothers treated with GCs during lactation (n = 18 litters, 79.6 ± 8% pups survived) and the lactation (time-matched) controls (n = 17 litters, 72.6 ± 9.5% pups survived, *z* = −0.63, *P* = 0.53).

Litter size did not differ between mothers treated with GCs during pregnancy or the controls at the first nest entry (Pregnancy GCs: n = 42 litters, 3.07 ± 0.14 pups, range = 1-5 pups; Pregnancy Controls: n = 31 litters, 2.9 ± 0.17 pups, range = 1-5 pups, effect of treatment, *t*52 = −0.19, *P* = 0.85) nor at the second nest entry (Pregnancy GCs: n = 23 litters, 2.75 ± 0.24 pups, range = 1-5 pups; Pregnancy Controls: n = 20 litters, 2.70 ± 0.16 pups, range = 1-4 pups, effect of treatment, *t*37 = 0.11, *P* = 0.92). Litter size also did not differ between mothers treated with GCs during lactation or the controls at the first nest entry (Lactation GCs: n = 18 litters, 2.67 ± 0.2 pups, range = 1-4 pups; Lactation Controls: n = 17 litters, 2.94 ± 0.13 pups, range = 2-4 pups, effect of treatment, *t*30 = −1.11, *P* = 0.27), nor at the second nest entry (Lactation GCs: n = 16 litters, 2.25 ± 0.2 pups, range = 1-3 pups; Lactation Controls: n = 14 litters, 2.57 ± 0.2 pups, range = 1-4 pup, effect of treatment, *t*25 = −1.31, *P* = 0.20).

The litter sex ratio (proportion of males) at the first nest entry did not differ between mothers treated with GCs during pregnancy or the controls (Pregnancy GCs: n = 42 litters, 53 ± 4% males; Pregnancy Controls: n = 29 litters, 40.9 ± 5% males, effect of treatment: *z* = 0.57, *P* = 0.57), nor at the second nest entry (Pregnancy GCs: n = 23 litters, 59.3 ± 7% males; Pregnancy Controls: n = 19 litters, 46.5 ± 7% males, effect of treatment: *z* = 0.26, *P* = 0.80). Similarly, the litter sex ratio at the first nest entry did not differ between mothers treated with GCs during lactation or the controls (Lactation GCs: n = 18 litters, 52.8 ± 7.2% males; Lactation Controls: n = 17 litters, 53.4 ± 7.9% males, effect of treatment: *z* = 0.09, *P* = 0.92) nor at the second nest entry (Lactation GCs: n = 16 litters, 58.3 ± 8.7% males; Lactation Controls: n = 14 litters, 64.9 ± 9.3% males, effect of treatment: *z* = −0.27, *P* = 0.79).

### Effects of treating pregnant females with GCs on offspring growth, oxidative stress, and telomere lengths

Offspring from mothers treated with GCs during pregnancy grew 17.0% faster (*t*_41.4_ = 3.07, *P* = 0.004, Table 2A, Fig. 1A) but were not larger in structural size (Table 2B, Fig. 1C), and did not differ in body condition (as reflected in their haematocrit levels: Table S2A) than those produced by control mothers. After correcting for the effects of age on the first body mass we recorded when pups were first weighed (“birth mass”) as well as other variables (Table 3A), offspring from mothers treated with GCs during pregnancy were 8.3% smaller than those from control mothers (*t*59.8 = −2.51, *P* = 0.015, Table 3A), suggesting that these pups exhibited catch-up growth after being born at a smaller body mass. There was no indication that the treatments had sex-specific effects on offspring growth or size (Table 2). There was also no indication of a trade-off between litter size and offspring growth or size during the years that we studied (Table 2) nor was there any evidence that treating mothers with GCs during pregnancy altered the relationship between litter size and growth rate, as indicated by the lack of significant interactions between treatment and litter size for offspring growth and size (Table 2). Because the treatments had no significant effects on litter size or litter sex ratio (see above), the effects of the treatments on offspring growth were not simply due to a reduction in litter size.

**Figure 1.**
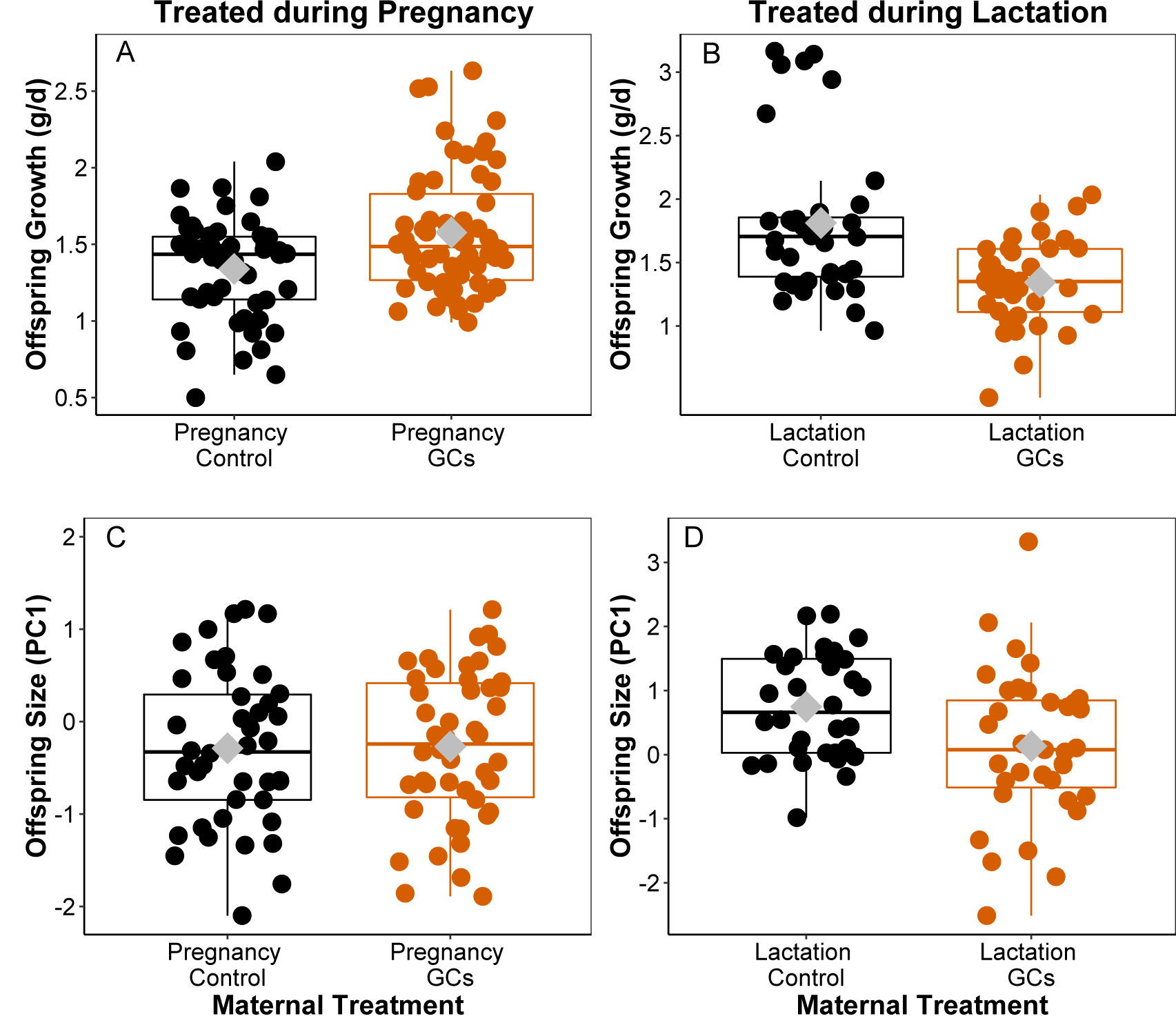
Effects of treating pregnant or lactating female red squirrels with glucocorticoids (GCs) on offspring postnatal growth and structural size. **A)** Offspring from females treated with GCs (n = 62 pups) during pregnancy grew significantly faster than those from controls (n = 52, *t*41.4 = 3.07, *P* = 0.004). **B)** Offspring from females treated with GCs during lactation (n = 36) grew significantly slower than those from controls (n = 36, *t*27 = –2.08, *P* = 0.047). **C & D)** Offspring from mothers treated with GCs either during pregnancy (n = 46) or lactation (n = 35) did not differ in structural size compared to those from controls (pregnancy: n = 42; lactation: n = 32, Tables 2 & 4). Offspring growth was measured as the change in body mass from ∼1 to ∼25 d of age. Offspring size is a composite variable where high scores of PC1 correspond to offspring with larger zygomatic arch widths and longer hind foot lengths when pups were ∼25 d of age. Results in Tables 2 & 4. Upper and lower hinges correspond to the first and third quartiles while white diamonds indicate means and horizontal line indicate medians.

**Table 2.** Effects of treating female red squirrels with glucocorticoids (GCs) during pregnancy on offspring postnatal growth (A) and structural size (B). Offspring growth is the linear change in body mass from ∼1 d to ∼25 d of age. Offspring size is a composite variable where high scores of PC1 correspond to offspring (one estimate of size obtained at ∼25 d of age) with larger zygomatic arch widths and longer hind foot lengths. Models contained random intercept term for litter identity (growth model: σ^2^ = 0.07; size model: σ^2^ = 0.29).

| (A) | Offspring trait | Variable | b | SE | *t* | df | *P*-value |
| --- | --- | --- | --- | --- | --- | --- | --- |
|  | **Growth** | **Intercept (2012, Control, Female)** | **1.81** | **0.13** | **14.2** | **35.6** | **<0.0001** |
|  |  | Year (2015) | -0.09 | 0.22 | -0.42 | 34.5 | 0.67 |
|  |  | **Year (2016)** | **-0.58** | **0.13** | **-4.3** | **34.2** | **<0.0001** |
|  |  | **Year (2017)** | **-0.74** | **0.13** | **-5.8** | **34.2** | **<0.0001** |
|  |  | **Sex (Male)** | **0.074** | **0.03** | **2.10** | **72.6** | **0.04** |
|  |  | Birth date | -0.001 | 0.05 | -0.03 | 34.6 | 0.98 |
|  |  | Litter size | 0.06 | 0.09 | 0.67 | 37.0 | 0.50 |
|  |  | **Treatment (GCs)** | **0.28** | **0.09** | **3.07** | **41.4** | **0.004** |
|  |  | Treatment (GCs) x Sex (Male) | -0.08 | 0.05 | -1.64 | 73.8 | 0.11 |
|  |  | Treatment (GCs) x Litter size | -0.08 | 0.11 | -0.68 | 36.7 | 0.50 |

Results based upon 114 offspring from 43 litters across 4 years
| (B) | Offspring trait | Variable | b | SE | *t* | df | *P*-value |
| --- | --- | --- | --- | --- | --- | --- | --- |
|  | **Size (PC1)** | **Intercept (2015, Control, Female)** | **0.14** | **0.41** | **0.35** | **31.9** | **0.72** |
|  |  | **Year (2016)** | **-1.14** | **0.43** | **-2.65** | **28.9** | **0.013** |
|  |  | Year (2017) | -0.17 | 0.43 | -0.39 | 29.4 | 0.70 |
|  |  | Sex (Male) | 0.27 | 0.16 | 1.69 | 59.6 | 0.09 |
|  |  | Birth date | 0.14 | 0.11 | 1.25 | 25.9 | 0.22 |
|  |  | Litter size | 0.02 | 0.21 | 0.11 | 33.2 | 0.92 |
|  |  | Treatment (GCs) | 0.10 | 0.25 | 0.39 | 44.8 | 0.70 |
|  |  | Treatment (GCs) x Sex (Male) | -0.10 | 0.22 | -0.42 | 60.8 | 0.67 |
|  |  | Treatment (GCs) x Litter size | -0.21 | 0.27 | -0.77 | 32.2 | 0.44 |

Results based upon 88 offspring from 34 litters across 3 years

**Table 3.**
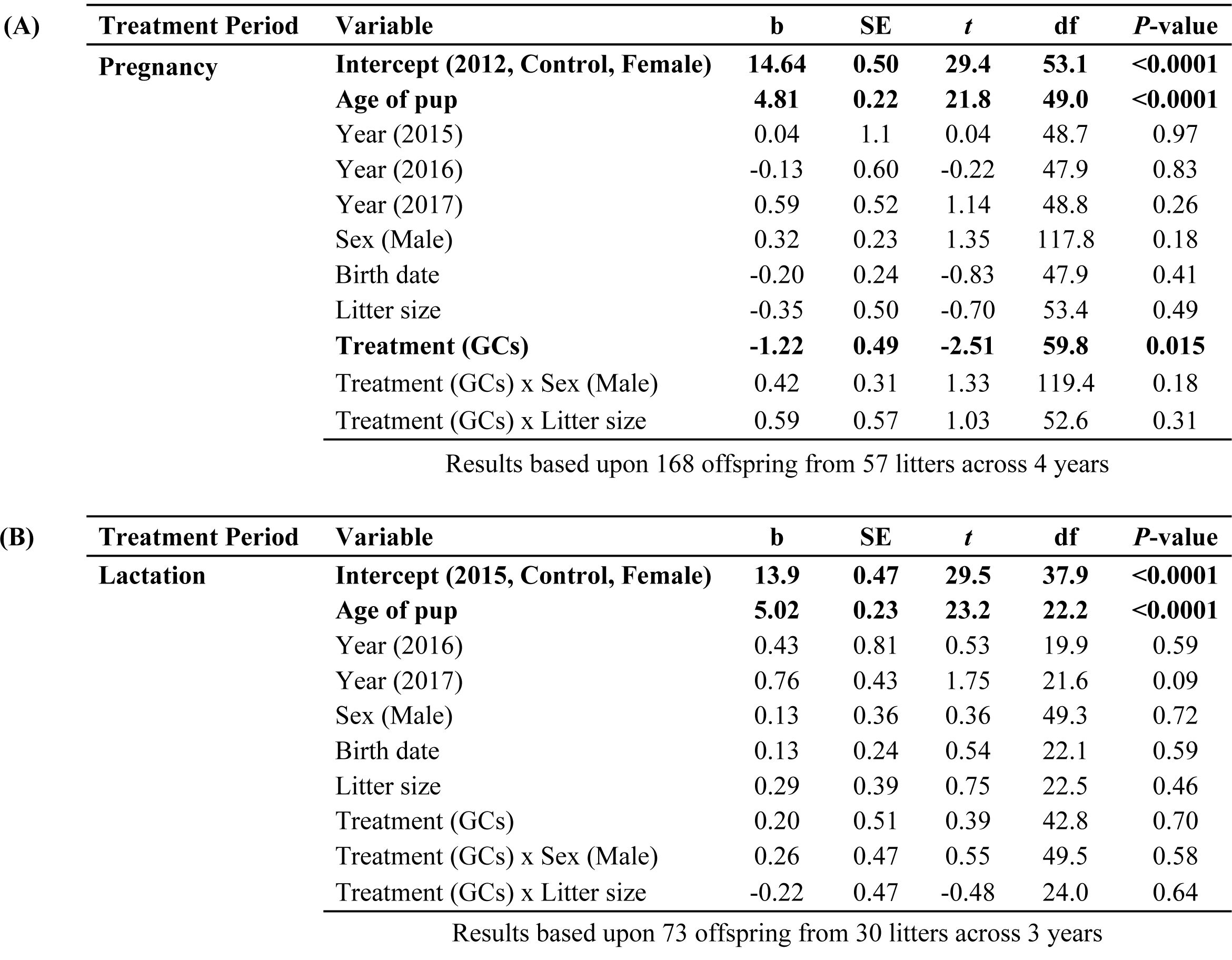
Effects of treating female red squirrels with different dosages of glucocorticoids (GCs) during (A) pregnancy or (B) lactation on the first body mass measure of offspring (“birth mass”). Offspring body mass was recorded soon after birth. Models contained random intercept term for litter identity (pregnancy model: σ^2^ = 2.3; lactation model: σ^2^ = 0.83).

| (A) | Treatment Period | Variable | b | SE | t | df | P-value |
| --- | --- | --- | --- | --- | --- | --- | --- |
|  | Pregnancy | Intercept (2012, Control, Female) | 14.64 | 0.50 | 29.4 | 53.1 | <0.0001 |
|  |  | Age of pup | 4.81 | 0.22 | 21.8 | 49.0 | <0.0001 |
|  |  | Year (2015) | 0.04 | 1.1 | 0.04 | 48.7 | 0.97 |
|  |  | Year (2016) | -0.13 | 0.60 | -0.22 | 47.9 | 0.83 |
|  |  | Year (2017) | 0.59 | 0.52 | 1.14 | 48.8 | 0.26 |
|  |  | Sex (Male) | 0.32 | 0.23 | 1.35 | 117.8 | 0.18 |
|  |  | Birth date | -0.20 | 0.24 | -0.83 | 47.9 | 0.41 |
|  |  | Litter size | -0.35 | 0.50 | -0.70 | 53.4 | 0.49 |
|  |  | Treatment (GCs) | -1.22 | 0.49 | -2.51 | 59.8 | 0.015 |
|  |  | Treatment (GCs) x Sex (Male) | 0.42 | 0.31 | 1.33 | 119.4 | 0.18 |
|  |  | Treatment (GCs) x Litter size | 0.59 | 0.57 | 1.03 | 52.6 | 0.31 |

Results based upon 168 offspring from 57 litters across 4 years
| (B) | Treatment Period | Variable | b | SE | t | df | P-value |
| --- | --- | --- | --- | --- | --- | --- | --- |
|  | Lactation | Intercept (2015, Control, Female) | 13.9 | 0.47 | 29.5 | 37.9 | <0.0001 |
|  |  | Age of pup | 5.02 | 0.23 | 23.2 | 22.2 | <0.0001 |
|  |  | Year (2016) | 0.43 | 0.81 | 0.53 | 19.9 | 0.59 |
|  |  | Year (2017) | 0.76 | 0.43 | 1.75 | 21.6 | 0.09 |
|  |  | Sex (Male) | 0.13 | 0.36 | 0.36 | 49.3 | 0.72 |
|  |  | Birth date | 0.13 | 0.24 | 0.54 | 22.1 | 0.59 |
|  |  | Litter size | 0.29 | 0.39 | 0.75 | 22.5 | 0.46 |
|  |  | Treatment (GCs) | 0.20 | 0.51 | 0.39 | 42.8 | 0.70 |
|  |  | Treatment (GCs) x Sex (Male) | 0.26 | 0.47 | 0.55 | 49.5 | 0.58 |
|  |  | Treatment (GCs) x Litter size | -0.22 | 0.47 | -0.48 | 24.0 | 0.64 |

Results based upon 73 offspring from 30 litters across 3 years

Offspring from mothers treated with GCs during pregnancy did not have higher oxidative stress levels in the blood, liver, or heart (Table S3, Fig. 2) and they also did not have shorter telomere lengths (Table S4, Fig. 3). Offspring from mothers treated with GCs during pregnancy or provided with supplemental food during pregnancy (controls) that grew faster did not have higher oxidative stress levels in blood, heart, or liver nor did they have shorter telomere lengths (Tables S3-S4). There was no indication that growth or its interaction with maternal treatment impacted oxidative stress levels (Table S3) or liver telomere lengths (Table S4) in offspring from females treated during pregnancy.

**Figure 2.**
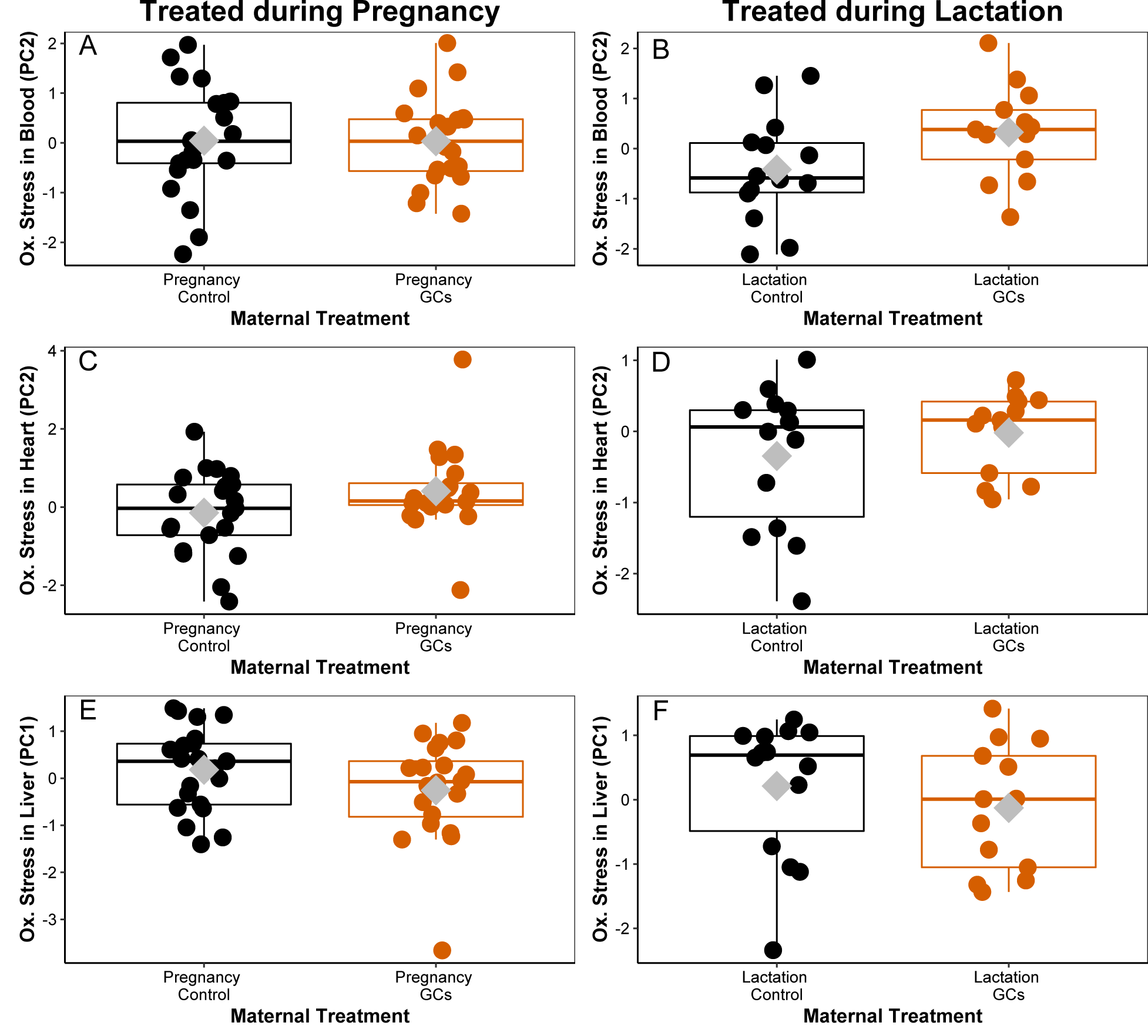
Effects of treating pregnant or lactating female red squirrels with glucocorticoids (GCs) on oxidative stress levels in blood, liver, and heart tissue from weaned juvenile red squirrels. There were no significant treatment effects on oxidative stress levels in the blood (**A, B**), heart (**C, D**), or liver (**E, F**) for offspring produced by mothers treated with GCs during pregnancy (n = 20 pups) or lactation (n = 13) compared with the controls (pregnancy: n = 21; lactation: n = 14, statistical results in Tables S3 & S5). Values on y-axes reflect a composite variable generated by separate principal component analyses for blood, heart, and liver tissue where high scores correspond to low levels of one or two of the antioxidants (TAC, SOD) and, for heart, higher levels of protein damage (PCC; see Table 1). Note differences between y-axes among the panels.

**Figure 3.**
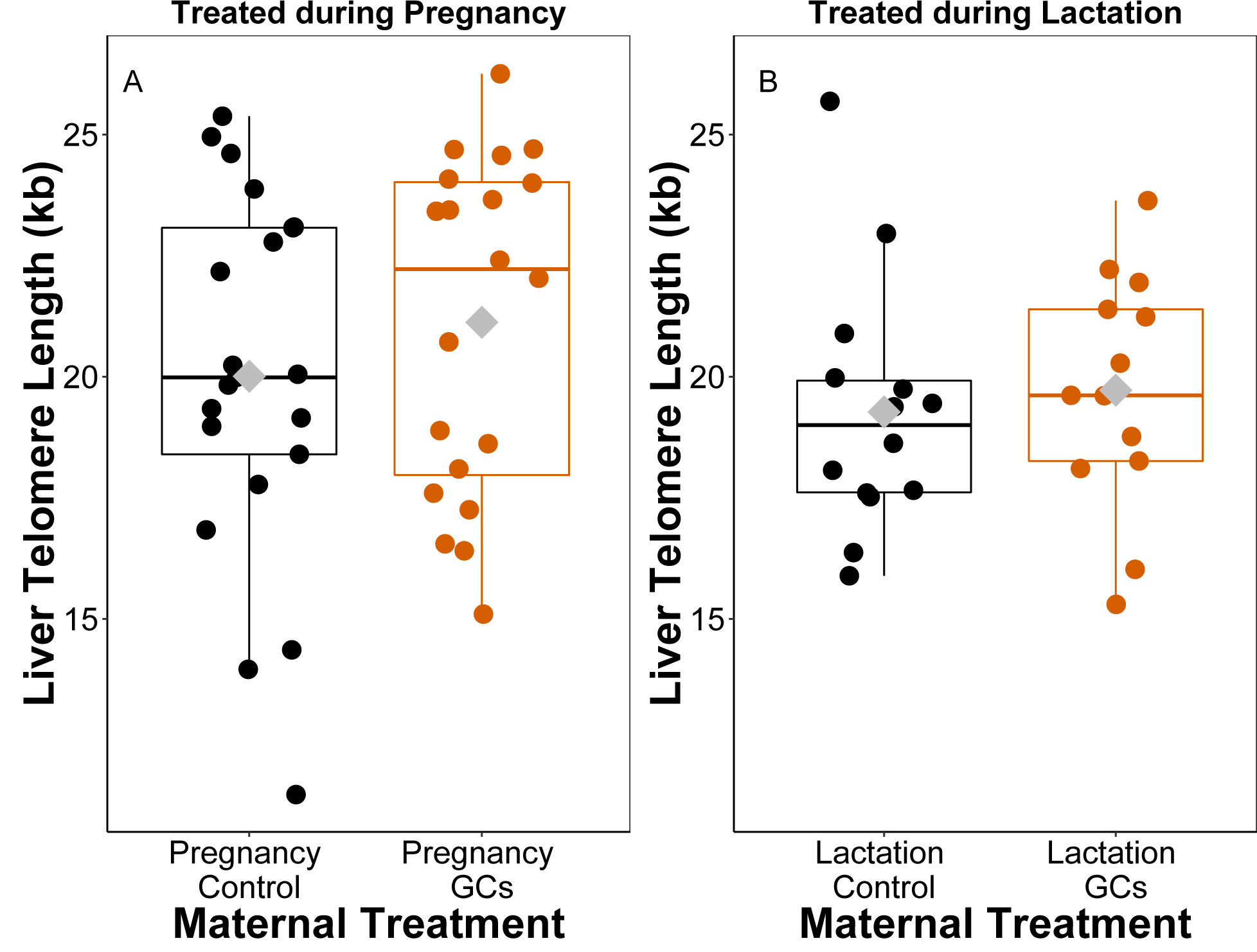
Effects of treating pregnant or lactating female red squirrels with glucocorticoids (GCs) on mean liver telomere lengths in weaned offspring. A & B) There were no significant treatment effects on liver telomere lengths for offspring produced by mothers treated with GCs during pregnancy (n = 20 pups) or lactation (n = 13) compared with the controls (pregnancy: n = 21; lactation: n = 14, statistical results in Table S4).

### Effects of treating lactating females with GCs on offspring growth, oxidative stress, and telomere lengths

Offspring from mothers treated with GCs during lactation grew 34.8% slower (*t*27 = – 2.08, *P* = 0.047, Table 4A, Fig. 1B), but were not significantly smaller in structural size (Table 4B, Fig. 1D), and did not differ in body condition (as reflected in haematocrit levels: Table S2B) or first body mass (Table 3B) from those of pups from control mothers. There was no indication that the treatments had sex-specific effects on offspring growth or size, as reflected in the lack of significant sex by treatment interactions (Table S4). There was also no indication of a trade-off between litter size and offspring growth or size (Table S4) and no evidence that treating females with GCs during lactation altered the relationship between litter size and growth rate, as indicated by the lack of significant interactions between treatment and litter size for offspring growth and size (Table S4). Because the treatments had no significant effects on litter size or litter sex ratio (see above), the effects of the treatments on offspring growth were not simply due to an increase in litter size.

**Table 4.**
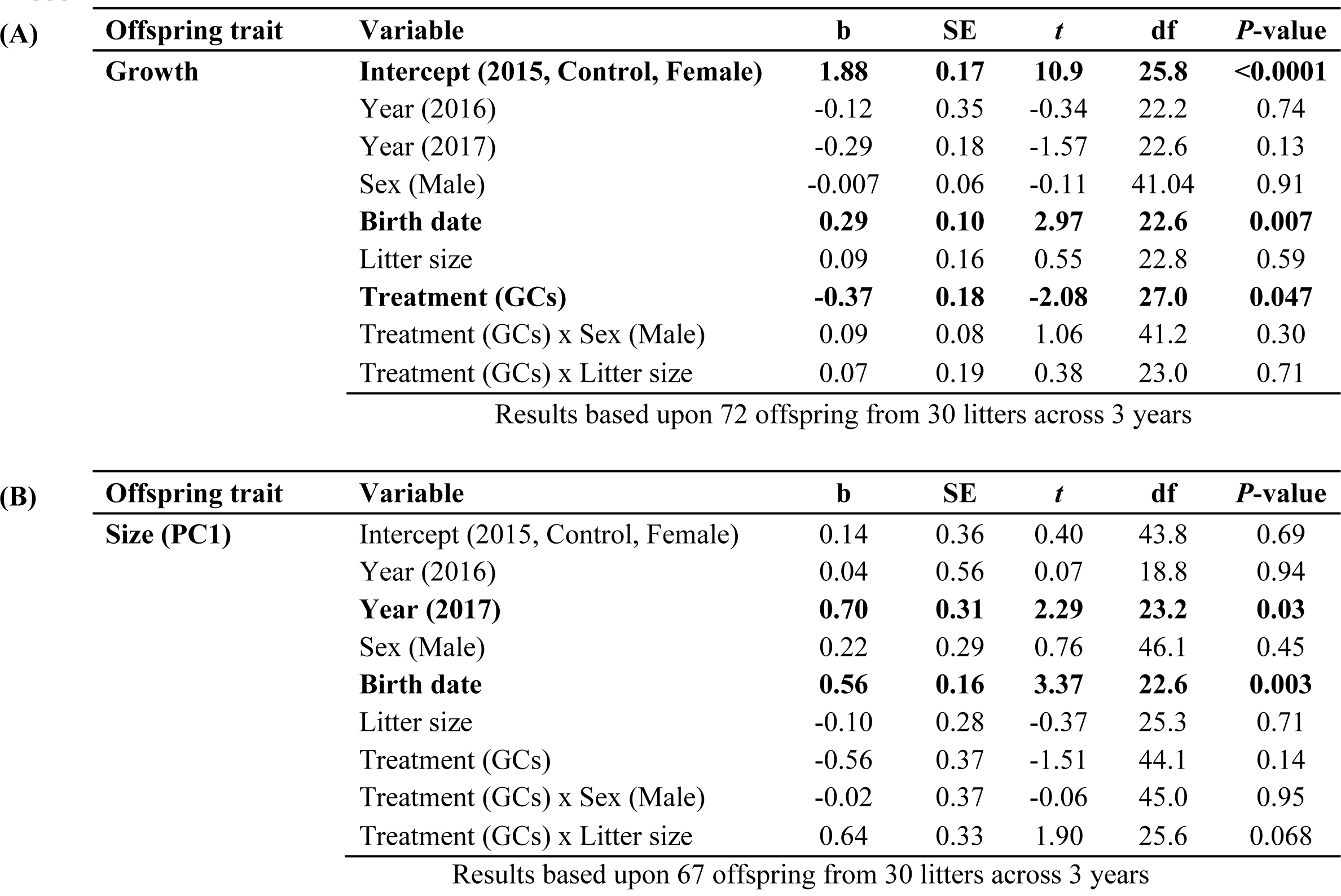
Effects of treating female red squirrels with GCs during lactation on offspring postnatal growth (A) and structural (skeletal) size (B). Offspring growth is the linear change in body mass from ∼1 d to ∼25 d of age. Offspring size is a composite variable where high scores of PC1 correspond to offspring (∼25 d of age) with larger zygomatic arch widths and longer hind foot lengths. Models contained random intercept term for litter identity (growth model: σ^2^ = 0.19; size model: σ^2^ = 0.69).

Offspring from mothers treated with GCs during lactation did not have higher oxidative stress levels than those from control mothers in any of the three tissues (blood, heart, liver: Table S5, Fig. 2B, 2D, 2F) and they also did not have shorter telomere lengths (Table S4, Fig. 3B). Offspring from mothers treated with GCs during lactation or provided with supplemental food during lactation (controls) that grew faster did not have higher oxidative stress levels in blood, heart, or liver nor did they have shorter telomere lengths (Table S4). There was also no indication that growth or its interaction with maternal treatment impacted oxidative stress levels (Table S5) or liver telomere lengths (Table S4) in offspring from females treated during lactation.

## Discussion

Mothers treated with GCs during pregnancy produced offspring that were lighter around birth but then exhibited significantly faster postnatal growth prior to weaning, whereas mothers treated with GCs during lactation tended to produce slower growing offspring that were of similar mass at birth. There were no treatment effects on offspring structural size, indicating that while offspring from mothers treated with GCs gained mass at a different rate than the controls, the treatments did not influence skeletal size. However, we only obtained one measure of structural size when offspring were ∼25 d of age, and therefore did not quantify any treatment effects in the rate of change in structural size as we did for body mass. Our results differ from a recent literature analysis across mammals showing that offspring from mothers experiencing late gestational stress grew more slowly before weaning (Berghänel et al., 2017). One explanation for the difference between this previous literature analysis and our results is that elevated GCs simply modulated the trade-off between offspring quantity and quality (e.g., producing small litters of fast-growing offspring) or ameliorated the trade-off between offspring quantity and quality (e.g., lessening the effect of increased litter size on the growth rate of each individual offspring: Stearns, 1992). However, this is unlikely because we found no treatment effects on litter size nor on the trade-off between litter size and growth rate. Thus, somehow females treated with GCs during pregnancy produced fast growing offspring without merely reducing their litter sizes, though it is notable that in the years in which we conducted this study, we also did not document a trade-off between litter size and offspring growth in any of the GC-treated or control females, as we have found previously (Dantzer et al., 2013). It is possible that we did not find evidence of a trade-off between litter size and offspring growth in the present study because both the GC-treated and control females were fed supplemental food and a previous study showed that *ad libitum* food-supplementation ameliorates this trade-off in red squirrels (Dantzer et al., 2013). GCs are metabolic hormones that have well-known effects on partitioning resources dedicated to survival vs. reproduction within-individuals (Wingfield et al., 1998). Here, it is possible that GCs also play a role in the partitioning of resources between mothers and offspring where they increase investment in offspring (increasing postnatal growth rates without a reduction in litter size) perhaps at the expense of maternal energy stores.

We did not find support for the hypothesis that elevated maternal GCs during pregnancy or lactation or increased offspring growth caused an elevation in oxidative stress levels or shortened telomere lengths in offspring. This differs from previous studies that have found that offspring growth is positively correlated with oxidative stress levels or inversely correlated with offspring telomere lengths (Haussmann and Heidinger, 2015; Monaghan and Hausssmann, 2015; Monaghan and Ozanne, 2018). This is surprising and requires explanation. First, we measured oxidative stress levels in offspring when they were weaned (∼70 d of age) whereas we treated their mothers with GCs either during pregnancy or early lactation. Thus, it is possible that the offspring in our study experienced elevated oxidative stress levels but these effects had disappeared by weaning. A second possibility is an artefact associated with selective disappearance of poor-quality individuals from those mothers treated with GCs during pregnancy, such as slow growing individuals with short telomeres dying before we could obtain our measures of oxidative stress and telomere lengths. This is unlikely as we observed no treatment effects on litter size or the reduction in litter size from the first to second nest entry. Finally, the fact that mothers treated with GCs did not produce offspring with elevated oxidative stress levels or shorter telomeres may have been because of the effects of maternally-derived GCs on offspring telomerase levels, an enzyme that is capable of rebuilding telomeres or buffering them from attrition (Blackburn, 2005). We did not measure telomerase levels, but a previous study showed that long-term exposure of laboratory rats to unpredictable stressors increased the production of telomerase (Beery et al., 2012). Thus, a testable prediction for future studies is that treating mothers with GCs promotes increases in telomerase or enzymatic antioxidant production that has protective effects on offspring. This would be consistent with predictions from the oxidative shielding hypothesis (Blount et al., 2016) that proposes that females may reduce their own levels of oxidative damage (perhaps by upregulation of enzymatic antioxidants) to mitigate their detrimental influence on offspring.

Our results indicate that elevated GCs can impact maternal investment in the current litter. Females experiencing elevated GCs during pregnancy increased their investment in the current litter (as indicated by an increase in postnatal growth) whereas females experiencing elevated GCs during lactation tended to reduce their investment in the current litter, the latter possibly being because mothers spent less time interacting or nursing their offspring, as in a previous study in laboratory rats (Nephew and Bridges, 2011). Alternatively, the reduction in offspring growth when mothers were treated with GCs during lactation could have been because milk quality or content was altered. Life history theory predicts that such changes in maternal investment in offspring could alter the survival or future reproduction of mothers (Stearns, 1992) or increase their oxidative stress levels (Monaghan and Haussmann, 2006; Blount et al., 2016). However, our previous work in red squirrels shows that their food caching nature results in unexpected patterns with respect to the potential costs of reproductive investment. Red squirrels can elevate reproductive output (i.e., producing a second litter or larger litters) in anticipation of increased future food abundance (Boutin et al., 2006), or produce faster growing offspring when the fitness payoffs warrant increased investment in the current litter (Dantzer et al., 2013). They seem to be able to do this without additional access to food except the food that they already have stored from the previous autumn (Humphries and Boutin, 2000; Boutin et al., 2006; Dantzer et al., 2013). Thus far, survival costs for this increased reproductive investment in the current litter exhibited by female red squirrels seem to be small or absent. Female red squirrels with increased reproductive effort do expend more energy (Humphries and Boutin, 2000; Fletcher et al., 2012) and experience increased oxidative protein damage (Fletcher et al., 2012) but we have not yet documented substantive survival costs for females that increase their reproductive output (Humphries and Boutin, 2000; Descamps et al., 2009). We have not yet quantified any oxidative or survival costs to mothers who were treated with GCs during pregnancy and who on average produced faster growing offspring. Unless females upregulate the production of protective enzymatic antioxidants or telomerase (Blount et al., 2016), it seems likely that females with elevated GCs would experience increased oxidative damage due to their elevated reproductive investment, or because of the elevated levels of GCs that they experience. For example, previous studies in red squirrels (Fletcher et al., 2012) and other species (Blount et al., 2016) highlight that increased reproductive investment or increased exposure to GCs (Kotrschal et al., 2007; You et al., 2009; Costantini et al., 2011) may elevate oxidative damage in breeding females with elevated GCs during reproduction.

Although it is not known whether increased antioxidants, reduced oxidative damage, or elongated telomeres actually cause an increase in longevity (Simons, 2015; but see Munoz-Lorente et al. 2019), our results suggest that fast growing offspring or those from mothers treated with GCs during pregnancy or lactation would not experience a reduction in lifespan. Our results and our previous study in red squirrels (Dantzer et al., 2013) show that maternal GCs during pregnancy or lactation can induce plasticity in offspring growth and that this plasticity should be adaptive for the fluctuating environments experienced by Yukon red squirrels (Dantzer et al., 2013). Specifically, when population density is elevated, females have elevated GCs and fast offspring growth increases offspring survival (Dantzer et al., 2013). We have shown here that females with elevated GCs during pregnancy produce faster growing offspring, whereas females with elevated GCs during lactation produce slower growing offspring. This suggests that plasticity in GC levels during female reproduction (e.g., females have elevated GCs during pregnancy but exert strong negative feedback on further GC production during lactation) would maximize female reproductive success when density is high. However, we did not find support for the hypothesis that elevated maternal GCs induce a faster pace of life where offspring grow faster and are more competitive early in life but this comes at some oxidative cost that may predict a shortened lifespan. Future studies should assess oxidative stress using an even broader array of measures than the few measures we used here, and will of course need to assess if elevated maternal GCs actually impact offspring lifespan. Our results to date indicate that the increases in maternal GCs during pregnancy in response to population density result in an adaptive maternal effect on offspring postnatal growth and do not pose a developmental constraint on offspring postnatal growth.

## Acknowledgements

We thank the Champagne and Aishihik First Nations and Agnes MacDonald for allowing us to study squirrels on their traditional territory. Special thanks to the field technicians that helped collect these data, especially to Monica Cooper, Stacie Evans, Zach Fogel, Claire Hoffmann, Noah Israel, Sean Konkolics, Laura Porter, Matt Sehrsweeney, Sam Sonnega, and Jess Steketee. This is publication 105 of the Kluane Red Squirrel Project.

## Data Availability

All data files are available on FigShare from the first author (https://figshare.com/account/home#/projects/71585). All other requests for data or analysis code will be fulfilled by the first author.

## Supplementary Results

**Table S1.** **Preliminary analyses for effects of treating female red squirrels with different dosages of glucocorticoids (GCs) during (A) pregnancy or (B) lactation on offspring postnatal growth.** Offspring growth is the linear change in body mass from ∼1 d to ∼25 d of age. The dosages of GCs are listed below with 0 mg (pregnancy control treatment, n = 52 pups; lactation control treatment, n = 36 pups) as the reference value. Models contained random intercept term for litter identity (pregnancy model: σ^2^ = 0.059; lactation model: σ^2^ = 0.23). Note that these results were used to justify our decision of lumping the different pregnancy GCs treatment groups (with different dosages of GCs) together.

| (A) | Treatment Period | Variable | b | SE | t | df | P-value |
| --- | --- | --- | --- | --- | --- | --- | --- |
|  | Pregnancy | Intercept (2012, Female, 0 mg) | **1.56** | **0.17** | **9.3** | **32.0** | **<0.0001** |
|  |  | Year (2015) | 0.06 | 0.21 | 0.31 | 32.2 | 0.76 |
|  |  | Year (2016) | -0.26 | 0.18 | -1.41 | 31.7 | 0.17 |
|  |  | **Year (2017)** | **-0.41** | **0.18** | **-2.28** | **31.6** | **0.03** |
|  |  | Sex (Male) | 0.037 | 0.02 | 1.45 | 74.5 | 0.15 |
|  |  | Birth date | -0.001 | 0.05 | -0.01 | 32.8 | 0.99 |
|  |  | Litter size | 0.035 | 0.05 | 0.72 | 34.2 | 0.48 |
|  |  | Treatment (GC dosage) |  |  |  |  |  |
|  |  | 3 mg GCs (n = 9 pups) | 0.37 | 0.24 | 1.52 | 30.9 | 0.14 |
|  |  | **6 mg GCs (n = 7 pups)** | **0.68** | **0.20** | **3.45** | **32.4** | **0.001** |
|  |  | 8 mg GCs (n = 45 pups) | 0.12 | 0.09 | 1.40 | 33.1 | 0.17 |
|  |  | **12 mg GCs (n = 1 pup)** | **0.91** | **0.31** | **2.95** | **36.9** | **0.005** |

Results based upon 114 offspring from 43 litters across 4 years
| (B) | Treatment Period | Variable | b | SE | t | df | P-value |
| --- | --- | --- | --- | --- | --- | --- | --- |
|  | Lactation | Intercept (2015, 0 mg, Female) | **1.96** | **0.19** | **9.9** | **23.7** | **<0.0001** |
|  |  | Year (2016) | -0.12 | 0.34 | -0.35 | 22.1 | 0.73 |
|  |  | Year (2017) | -0.49 | 0.25 | -1.94 | 22.5 | 0.065 |
|  |  | Sex (Male) | 0.04 | 0.04 | 1.10 | 42.5 | 0.27 |
|  |  | **Birth date** | **0.25** | **0.10** | **2.46** | **22.8** | **0.021** |
|  |  | Litter size | 0.15 | 0.09 | 1.53 | 23.1 | 0.14 |
|  |  | Treatment (GCs) |  |  |  |  |  |
|  |  | 8 mg GCs (n = 24 pups) | -0.16 | 0.22 | -0.74 | 22.1 | 0.47 |
|  |  | 12 mg GCs (n = 13 pups) | -0.55 | 0.27 | -1.99 | 23.1 | 0.058 |

Results based upon 72 offspring from 30 litters across 3 years

**Table S2.**
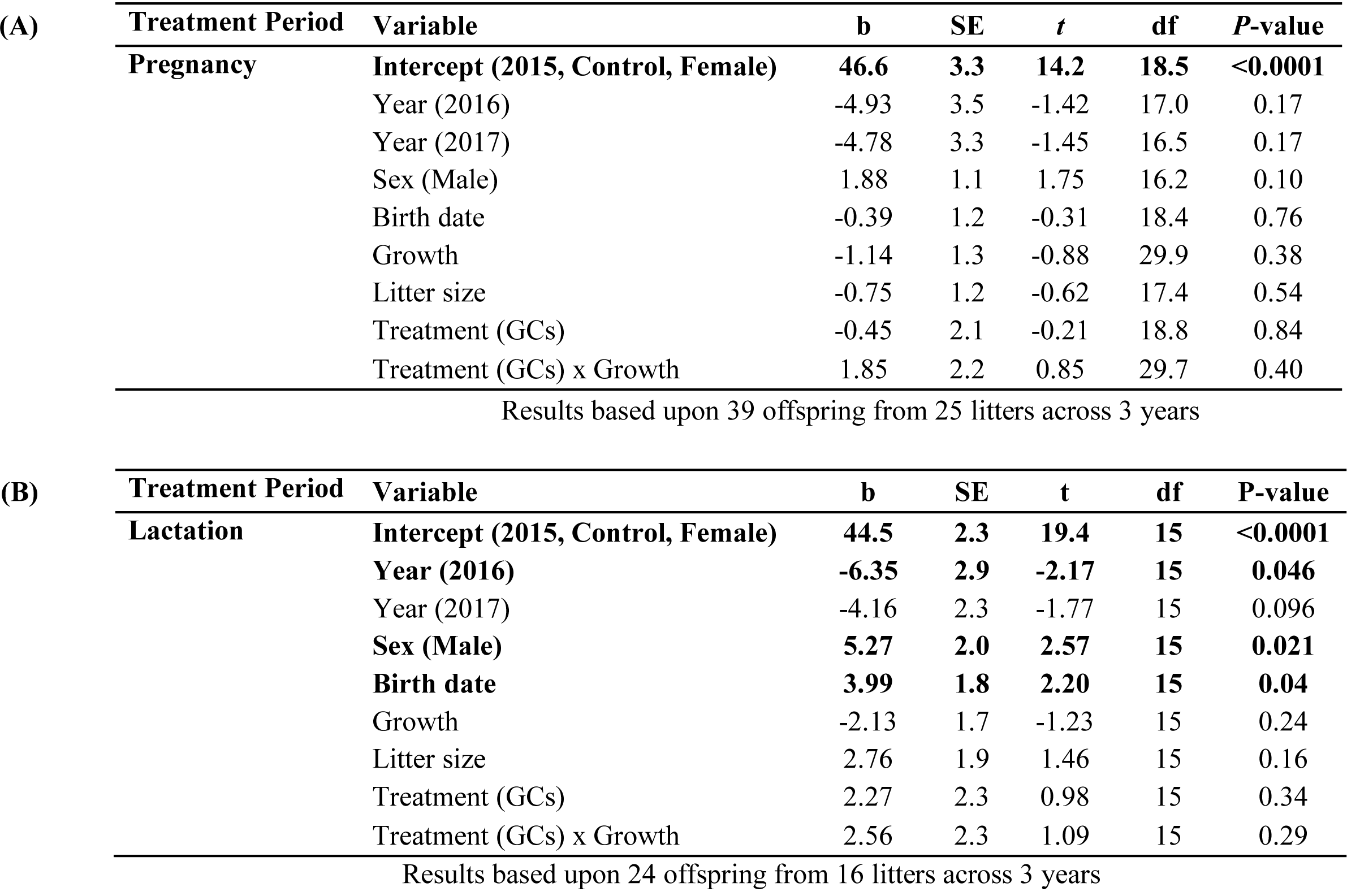
**Effects of treating female red squirrels with GCs during (A) pregnancy or (B) lactation on offspring haematocrit levels (packed red blood cell volume) collected from weaned offspring**. Results for the pregnancy model contained random intercept term for litter identity (pregnancy: σ^2^ = 18.8) whereas results for the lactation model are from a general linear model.

**Table S3.**
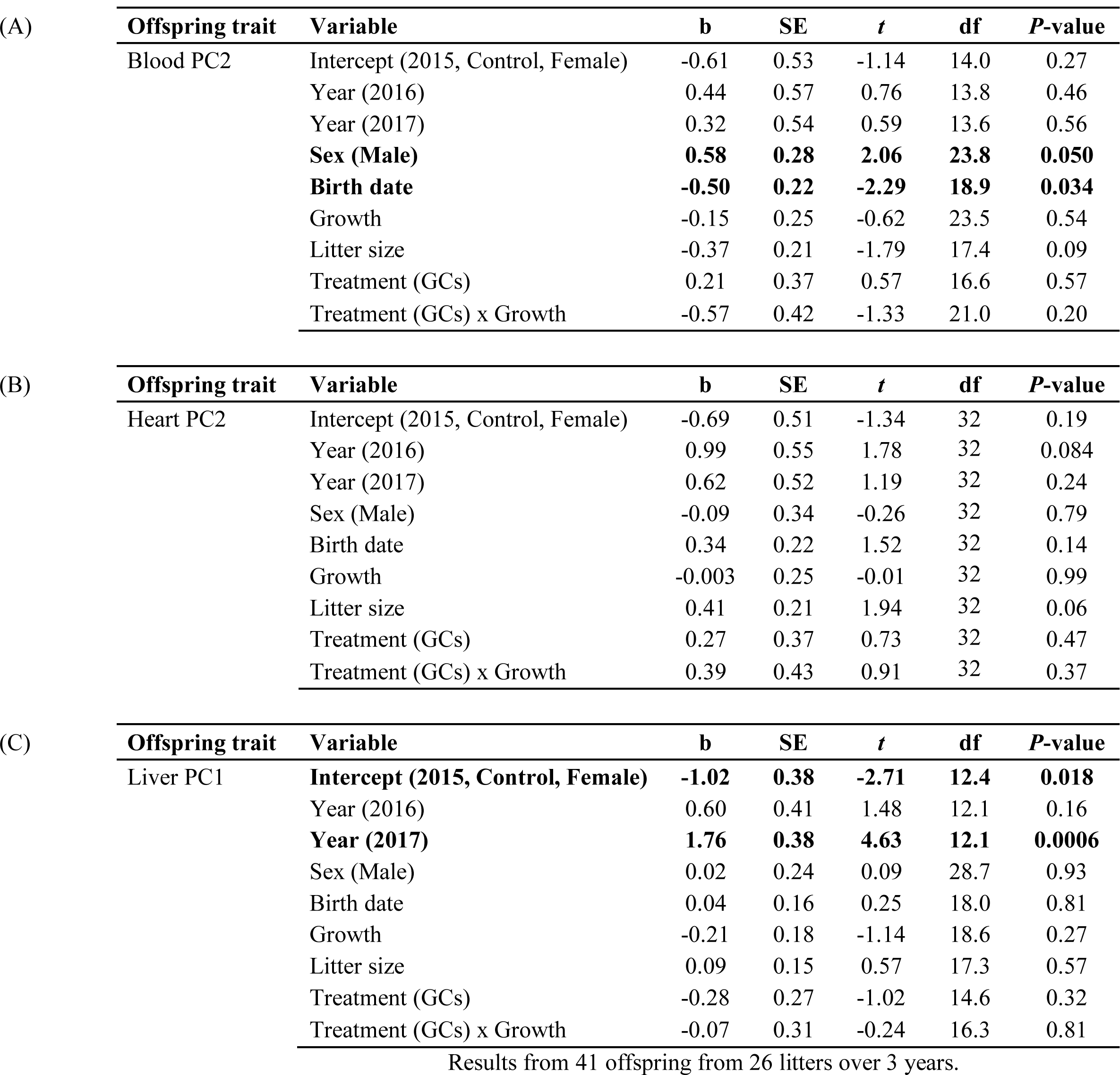
**Effects of treating pregnant red squirrels with GCs on oxidative stress levels in (A) blood, (B) heart, and (C) liver tissue from weaned offspring.** High PC scores correspond to low levels of antioxidants and, for heart, higher levels of protein damage (see Table 1). Models for blood and liver tissues contained random intercept term for litter ID (blood: σ^2^ = 0.32; liver: σ^2^ = 0.04).

**Table S4.** **Effects of treating female red squirrels with GCs during (A) pregnancy or (B) lactation on liver telomere lengths (kb) of weaned offspring.** Telomeres measured in DNA from liver tissue using the TRF method. Models contained random intercept term for litter identity (pregnancy: σ^2^ = 0.96; lactation: σ^2^ = 6.7).

| **Treatment Period** | **Variable** | **b** | **SE** | **t** | **df** | **P-value** |
| --- | --- | --- | --- | --- | --- | --- |
| **Pregnancy** | **Intercept (2015, Control, Female)** | **20.59** | **1.40** | **14.7** | **15.5** | **<0.0001** |
|  | **Year (2016)** | **3.69** | **1.41** | **2.61** | **13.0** | **0.022** |
|  | Year (2017) | -3.07 | 1.63 | -1.88 | 19.0 | 0.076 |
|  | Sex (Male) | 0.33 | 0.76 | 0.43 | 26.4 | 0.67 |
|  | Birth date | 0.54 | 0.54 | 1.01 | 18.2 | 0.32 |
|  | Growth | -0.30 | 0.62 | -0.49 | 20.7 | 0.63 |
|  | Litter size | -0.39 | 0.51 | -0.76 | 17.7 | 0.45 |
|  | Liver PC1 | 0.23 | 0.57 | 0.41 | 30.8 | 0.69 |
|  | Treatment (GCs) | 0.42 | 0.92 | 0.45 | 15.5 | 0.65 |
|  | Treatment (GCs) x Growth | 1.18 | 1.04 | 1.14 | 17.5 | 0.27 |

Results based upon 41 offspring from 26 litters across 3 years
| **Treatment Period** | **Variable** | **b** | **SE** | **t** | **df** | **P-value** |
| --- | --- | --- | --- | --- | --- | --- |
| **Lactation** | **Intercept (2015, Control, Female)** | **19.03** | **1.97** | **9.67** | **14.9** | **<0.0001** |
|  | Year (2016) | 2.88 | 2.61 | 1.10 | 11.5 | 0.29 |
|  | Year (2017) | 0.29 | 2.68 | 0.11 | 15.8 | 0.91 |
|  | Sex (Male) | 0.12 | 1.04 | 0.11 | 11.1 | 0.91 |
|  | Birth date | -0.09 | 1.24 | -0.07 | 12.1 | 0.94 |
|  | Growth | -0.08 | 0.86 | -0.09 | 16.6 | 0.93 |
|  | Litter size | 0.57 | 1.52 | 0.37 | 11.9 | 0.71 |
|  | Liver PC1 | -0.41 | 0.85 | -0.48 | 16.2 | 0.64 |
|  | Treatment (GCs) | 0.52 | 1.49 | 0.35 | 11.3 | 0.73 |
|  | Treatment (GCs) x Growth | 1.03 | 1.17 | 0.88 | 14.1 | 0.39 |

Results based upon 27 offspring from 18 litters across 3 years

**Table S5.** **Effects of treating lactating red squirrels with GCs on oxidative stress levels in (A) blood, (B) heart, and (C) liver tissue from weaned offspring.** High PC scores correspond to low levels of antioxidants and, in heart tissue, higher levels of protein damage (Table 1). The model for liver contained a random intercept term for litter identity (liver: σ2 = 0.33).

| (A) | Offspring trait | Variable | b | SE | t | df | P-value |
| --- | --- | --- | --- | --- | --- | --- | --- |
|  | Blood PC2 | Intercept (2015, Control, Female) | 0.21 | 0.52 | 0.40 | 18 | 0.69 |
|  |  | Year (2016) | -0.16 | 0.73 | -0.22 | 18 | 0.83 |
|  |  | Year (2017) | -0.56 | 0.57 | -0.99 | 18 | 0.34 |
|  |  | Sex (Male) | -0.53 | 0.49 | -1.06 | 18 | 0.30 |
|  |  | Birth date | -0.23 | 0.39 | -0.59 | 18 | 0.56 |
|  |  | Growth | 0.39 | 0.33 | 1.17 | 18 | 0.26 |
|  |  | Litter size | -0.42 | 0.46 | -0.92 | 18 | 0.37 |
|  |  | Treatment (GCs) | 0.44 | 0.44 | 1.01 | 18 | 0.32 |
|  |  | Treatment (GCs) x Growth | -0.29 | 0.47 | -0.62 | 18 | 0.54 |

| (B) | Offspring trait | Variable | b | SE | t | df | P-value |
| --- | --- | --- | --- | --- | --- | --- | --- |
|  | Heart PC2 | Intercept (2015, Control, Female) | **-1.14** | **0.41** | **-2.78** | **18** | **0.012** |
|  |  | Year (2016) | 0.55 | 0.57 | 0.97 | 18 | 0.35 |
|  |  | Year (2017) | 0.84 | 0.44 | 1.87 | 18 | 0.08 |
|  |  | Sex (Male) | 0.36 | 0.39 | 0.92 | 18 | 0.37 |
|  |  | Birth date | 0.23 | 0.31 | 0.76 | 18 | 0.46 |
|  |  | Growth | -0.09 | 0.26 | -0.36 | 18 | 0.72 |
|  |  | Litter size | 0.27 | 0.36 | 0.74 | 18 | 0.47 |
|  |  | Treatment (GCs) | 0.58 | 0.34 | 1.68 | 18 | 0.11 |
|  |  | Treatment (GCs) x Growth | -0.16 | 0.37 | -0.44 | 18 | 0.66 |

| (C) | Offspring trait | Variable | b | SE | t | df | P-value |
| --- | --- | --- | --- | --- | --- | --- | --- |
|  | Liver PC1 | Intercept (2015, Control, Female) | **-1.35** | **0.40** | **-3.38** | **11.9** | **0.005** |
|  |  | Year (2016) | 0.71 | 0.62 | 1.14 | 9.6 | 0.28 |
|  |  | **Year (2017)** | **2.24** | **0.47** | **4.81** | **9.9** | **0.0007** |
|  |  | Sex (Male) | 0.12 | 0.31 | 0.40 | 14.1 | 0.69 |
|  |  | Birth date | -0.18 | 0.31 | -0.59 | 11.4 | 0.57 |
|  |  | Growth | -0.29 | 0.23 | -1.25 | 17.6 | 0.22 |
|  |  | Litter size | -0.35 | 0.37 | -0.93 | 10.7 | 0.37 |
|  |  | Treatment (GCs) | -0.12 | 0.37 | -0.32 | 10.1 | 0.75 |
|  |  | Treatment (GCs) x Growth | -0.13 | 0.33 | -0.40 | 17.9 | 0.69 |

Results from 27 offspring from 18 litters over 3 years.

